# Collided ribosomes are rescued by the endonuclease Rae1 and *trans-*translation

**DOI:** 10.64898/2026.09.23.753862

**Authors:** Daniel D. Tetreault, Katrina Callan, Cassidy R. Prince, Heather A. Feaga

## Abstract

Ribosome stalling is a major problem in all domains of life. When a ribosome stalls, trailing ribosomes may catch up to and collide with the stalled ribosome, depleting protein synthesis capacity. Here, we describe a novel pathway used by Gram-positive bacteria to rescue ribosome collisions. We used the ATPase defective ABCF protein YdiF(EQ_2_) to induce ribosome stalling and collisions in *Bacillus subtilis*. Ribosome profiling (Ribo-seq) of YdiF(EQ_2_)-expressing cells revealed that collided ribosomes are enriched for tmRNA, a functional RNA involved in *trans*-translation. We confirmed that tmRNA tagging activity is globally increased upon expression of any ATPase defective ABCF as well as in cells treated with the collision-inducing antibiotic erythromycin, suggesting this is a generalizable mechanism to rescue stalled and collided ribosomes. The global increase in tmRNA tagging that occurred in response to both erythromycin and YdiF(EQ2) induced collisions was dependent on the Rae1 endonuclease. Loss of *trans*-translation in cells experiencing widespread ribosome collisions leads to a severe fitness defect, consistent with the importance of this pathway in rescuing ribosomes stalled on truncated mRNAs that result from ribosome collisions. Altogether, our work supports a model in which Rae1 cleaves mRNA on collided ribosomes, thereby generating a truncated mRNA substrate for *trans*-translation and leading to rescue and recycling of the collided ribosomes. We term this mechanism <u>C</u>ollision-<u>A</u>ssociated <u>R</u>ae1-induced *trans-*Translation (CART). CART broadens the repertoire of tools that bacteria use to manage ribosome collisions.

**Significance:** Prolonged ribosome stalling leads to ribosome collisions, which are rescued by specialized factors. While ribosome collisions have been extensively studied in eukaryotes, our understanding of collision rescue in bacteria is in its infancy. Data described here are the first to directly show that tmRNA mediates rescue of collided ribosomes in a Gram-positive bacterium, *Bacillus subtilis*. This pathway is analogous to what occurs in model organisms such as *Escherichia coli* and *Saccharomyces cerevisiae*, but relies on an unrelated nuclease, Rae1. Since *B. subtilis* and *E. coli* are on opposite ends of the bacterial phylogenetic tree, and since Rae1 is broadly conserved in bacteria, our findings suggest that mRNA cleavage arose convergently in distantly related bacteria as a strategy to rescue ribosome collisions. Convergent evolution of these pathways highlights the importance of rescuing collided ribosomes in all organisms. Moreover, insights into ribosome rescue in *E. coli* and *B. subtilis* can guide studies of ribosome rescue in bacteria with intermediary phylogenetic relatedness to these two model organisms.

## Introduction

Ribosomes frequently become stalled at difficult-to-translate sequences, such as polyproline tracts, poly-basic or poly-acidic tracts, or tracts of rare codons (1–6). Since ribosomes are loaded sequentially onto mRNA, ribosome stalling causes trailing ribosomes to catch up and collide with the leading ribosome, forming a collided disome (7,8). Collided ribosomes require specialized rescue factors. In *Bacillus subtilis*, ribosome collisions are rescued by a split-and-clear pathway (9–16). First, the ATPase MutS2 binds at the disome interface and splits the large and small ribosomal subunits of the stalled ribosome from the mRNA (13,14). Ribosome splitting leaves the large subunit obstructed with peptidyl-tRNA. RqcH, along with either RqcP or YlmH, catalyzes non-templated addition of alanine to the peptide trapped in the large subunit (10,11,15). Alanine-tailing at the C-terminus of the nascent peptide allows it to slip out of the exit tunnel, exposing the ester linkage between the peptide and tRNA to the cytosol so that PTH can cleave the nascent peptide from the tRNA, clearing the obstructed large subunit (17). The truncated alanine-tagged peptide is then targeted for degradation by proteases (9).

Another strategy for rescuing ribosome collisions is the cut-and-rescue pathway used by *Escherichia coli* (16,18). Here, the endonuclease SmrB recognizes the collision interface and cleaves the mRNA on the collided ribosomes (18). Truncated mRNA on the trailing ribosome allows this ribosome to be a substrate for *trans-*translation, a pathway that rescues ribosomes trapped on mRNAs lacking stop codons (19–23). *trans-*Translation is mediated by tmRNA, a bifunctional RNA containing a tRNA-like domain charged with alanine and an mRNA-like domain that feeds the mRNA channel an intact open reading frame. The tmRNA ORF provides an in-frame stop codon and encodes a degron tag that targets the truncated nascent peptide for degradation (24–26). Importantly, MutS2 in *B. subtilis* is also an SMR domain-containing protein, and while both SmrB and MutS2 use their SMR domains to bind small subunit proteins at the collision interface, only SmrB cleaves the associated mRNA (13,14).

Cells encode numerous pathways to prevent ribosome stalling and subsequent ribosome collisions. During elongation, the ribosomal E site releases deacylated tRNAs, but when the ribosome stalls, the E site is left temporarily unoccupied. Recent work has highlighted a novel class of ribosome-associated proteins that bind the E site and resolve ribosome stalling: ABCF ATPases (27–35). ABCF ATPases are a distinct subfamily of ATP-binding cassette (ABC) ATPases that share a conserved domain arrangement including an arm domain that interacts with E-site proteins, a P-site tRNA interaction motif (PtIM) that contacts the P-site tRNA, and two ATP-binding cassettes (**Fig. 1A**) (29). ATP hydrolysis is thought to promote ABCF release from the ribosome, since inactivation of the ATP hydrolysis activity through mutation of the Walker B motif in both ATPase cassettes (EQ_2_ mutation) increases affinity for the ribosome and traps the ribosome in a state that prevents translocation (29,30). Expression of any ABCF(EQ_2_) mutant in *E. coli* globally inhibits protein synthesis, presumably through widespread ribosome stalling (30,31).

**Figure 1.**
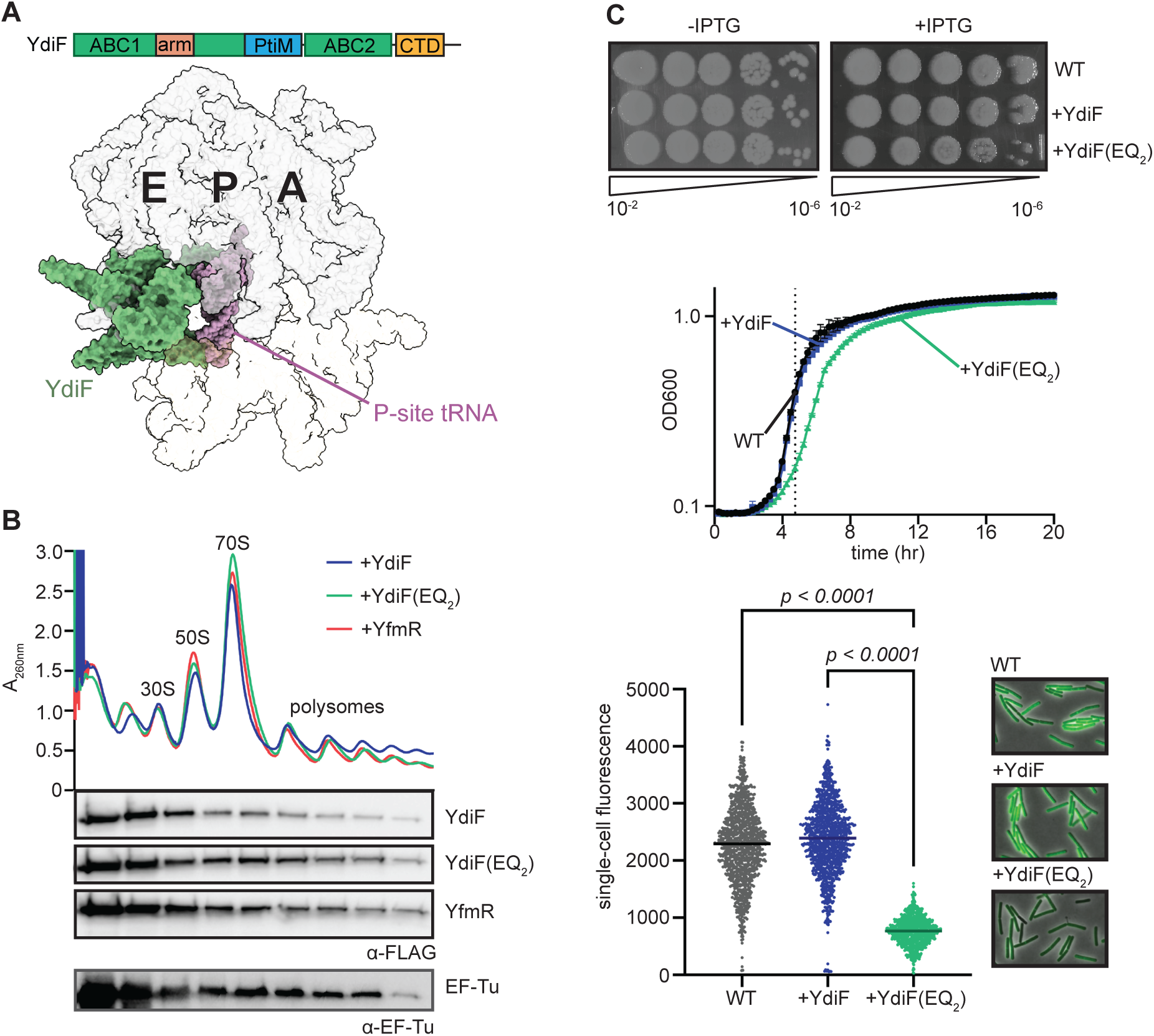
Inactive ABCF ATPase YdiF decreases cell growth and global translation. **(A)** Schematic showing the domain architecture of YdiF, including its ATP-binding cassettes, arm, P-site tRNA interacting motif (PtIM), and C-terminal domain extension. Alphafold predicted structure of *B. subtilis* YdiF bound to the ribosomal E-site via overlay atop *E. coli* EttA (PDB 3J5S, (88)). **(B)** Sucrose density gradients and western blot lysates harvested from cells expressing YdiF, YdiF(EQ_2_), and YfmR. YfmR and EF-Tu are shown as a positive control for ribosome association. **(C)** YdiF or YdiF(EQ_2_) expressing cells were grown on solid agar or in liquid media at 37°C. Protein synthesis of these cells grown in liquid was determined using BONCAT. The dotted line on the growth curve indicates the time point at which cells were collected for BONCAT. Growth curve error bars represent the SEM. Data points on graph of BONCAT experiment each represent the mean fluorescence of a single imaged cell. Representative microscope images from BONCAT labeling are shown at right. BONCAT *p-*values are the result of one-way ANOVA and Tukey’s multiple comparisons test.

Here, we used two approaches to cause widespread ribosome collisions in *B. subtilis* and determine how collisions are rescued. First, we used an ATP hydrolysis deficient ABCF (YdiF(EQ_2_)) as an inducible genetic tool to stall ribosomes and verified that this approach generates ribosome collisions using sucrose density gradient ultracentrifugation. Second, we used the antibiotic erythromycin, which causes ribosome stalling at specific motifs and is therefore a common tool in the field for generating ribosome collisions (36–38). Both approaches increased tmRNA-mediated tagging in the cells, which was dependent on the endonuclease Rae1. Moreover, ribosome profiling of cells expressing YdiF(EQ_2_) revealed that collided ribosomes exhibit increased occupancy on tmRNA. Altogether, these findings support a model in which Rae1 cleaves mRNA on collided ribosomes, thereby generating a truncated mRNA that allows tmRNA to rescue the ribosome. This pathway expands the repertoire of mechanisms that *B. subtilis* uses to rescue collided ribosomes.

## Results

### Inactive ABCF ATPase YdiF decreases translation

To identify mechanisms of ribosome rescue *in vivo*, we constructed a genetic tool to induce ribosome collisions in *B. subtilis* using an ATPase defective version of the YdiF ABCF ATPase. We mutated glutamate to glutamine (EQ_2_) in both Walker B domains of YdiF (39). To confirm that wild-type YdiF and YdiF(EQ_2_) associate with actively translating ribosomes, we resolved the ribosomes of cells expressing YdiF-FLAG and YdiF(EQ_2_)-FLAG using sucrose density gradient ultracentrifugation. As a positive control for ribosome association, we used the previously characterized *B. subtilis* ABCF ATPase, YfmR (32,33). We detected both YdiF-FLAG and YdiF(EQ_2_)-FLAG in dense polysome fractions (**Fig. 1B**). This binding pattern is comparable to that of YfmR and to the canonical elongation factor EF-Tu, which binds the ribosome at each cycle of elongation. These results indicate that YdiF-FLAG and YdiF(EQ_2_)-FLAG strongly associate with actively translating ribosomes.

Next, we determined the effect of YdiF(EQ_2_) expression on cell growth and translation. We prepared serial dilutions of cells expressing IPTG-inducible YdiF or YdiF(EQ_2_) and spotted them on plates with and without IPTG. We also compared growth in liquid media. Under both conditions, YdiF-expressing cells grew comparably to wild-type cells while YdiF(EQ_2_)-expressing cells exhibited a modest growth defect (**Fig. 1C**). To test whether expression of YdiF(EQ_2_) interfered with global protein synthesis, we performed bio-orthogonal noncanonical amino acid tagging (BONCAT) in wild-type cells and in cells expressing YdiF or YdiF(EQ_2_). BONCAT enables quantification of *de novo* protein synthesis with single-cell resolution via the incorporation of a clickable methionine analogue (HPG), which is then covalently linked to a fluorophore to measure the amount of HPG incorporation into nascent proteins (40). Cells were collected at mid-log phase at the time-point indicated by the dashed line (**Fig. 1C**). As expected, YdiF overexpression did not impact global translation levels, which remained comparable to global translation levels in wild-type cells (**Fig. 1C**). However, YdiF(EQ_2_) expression caused a significant decrease in global translation throughout the cell population. Representative fields of view used to quantify HPG incorporation reveal a visible decrease in fluorescence in cells expressing YdiF(EQ_2_) compared to both wild-type cells and cells expressing YdiF (**Fig. 1C**). These data suggest that expression of YdiF(EQ_2_) incurs a modest growth defect via interfering with global protein synthesis.

### Expression of YdiF(EQ_2_) generates ribosome collisions

YdiF(EQ_2_) binds tightly to the ribosome and globally reduces protein synthesis (**Fig. 1B, 1C**). To determine whether YdiF(EQ_2_) induces ribosome collisions, we used MNase digestion to test for the presence of nuclease-resistant disomes in cells with and without YdiF(EQ_2_) expression. When ribosomes are spaced along mRNAs, the nuclease MNase can access and cleave mRNA between them, thereby collapsing polysomes to monosomes (41). In contrast, collided ribosomes occlude MNase, preventing polysome collapse, and these species appear as disomes and trisomes on sucrose density gradients. Therefore, nuclease digestion is commonly used to assay levels of ribosome collisions (14,18,41–43).

When we treated ribosomes pelleted from wild-type cells with MNase, most polysomes collapsed into a single monosome peak (**Fig. 2A**). In contrast, when we digested ribosomes pelleted from cells expressing YdiF(EQ_2_), there were high levels of denser nuclease-resistant disome and trisome species at depths of 33 mm and 40 mm in the sucrose gradient, respectively (**Fig. 2A**). The intensity of the monosome peak obtained through polysome collapse correspondingly decreased in cells expressing YdiF(EQ_2_), consistent with MNase being unable to access mRNA between collided ribosomes. These data demonstrate that overexpression of YdiF(EQ_2_) generates ribosome collisions. Widespread ribosome collisions deplete the pool of available ribosomes, which likely explains the reduced protein synthesis and growth rate we observed in liquid culture (**Fig. 1C**).

**Figure 2.**
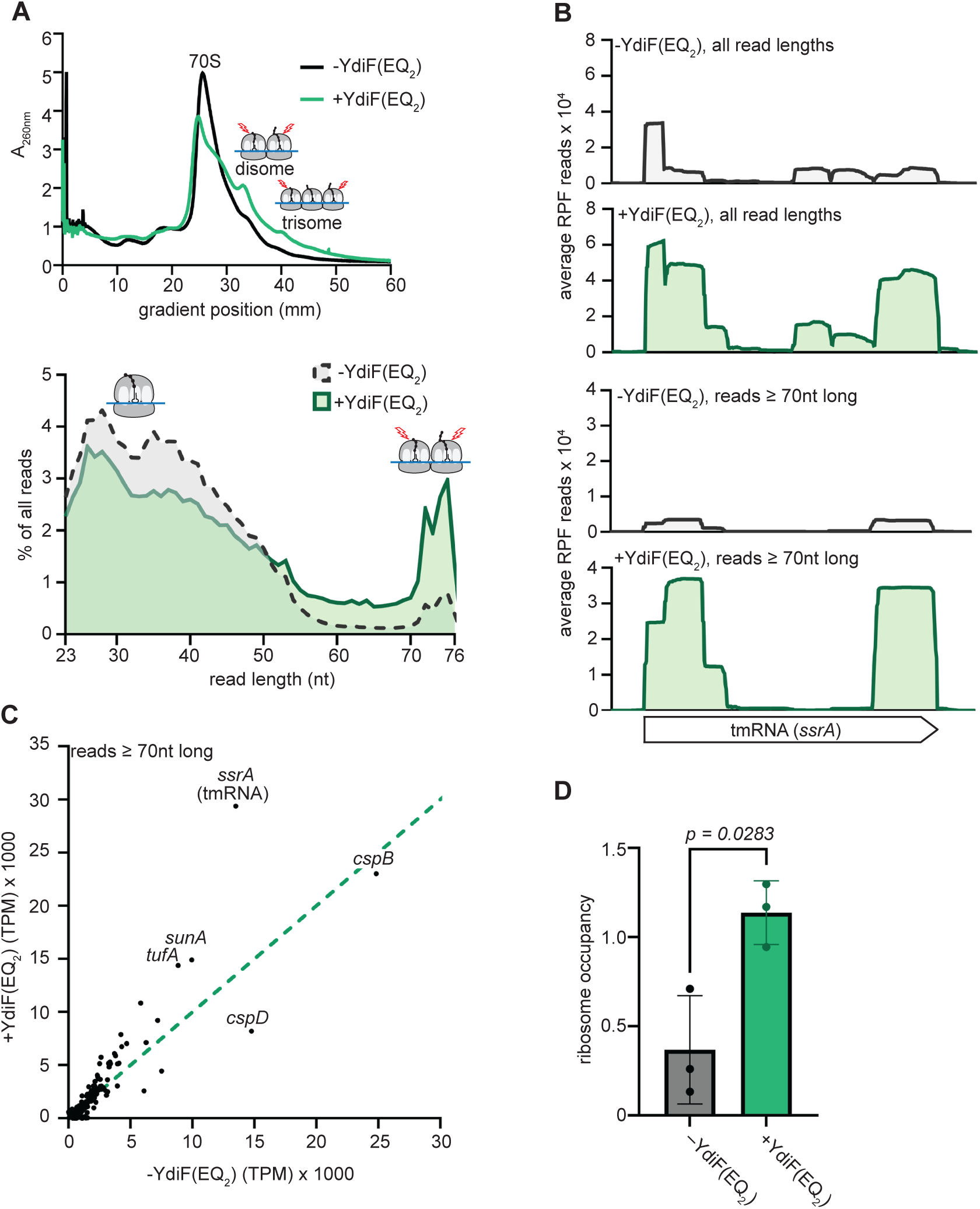
Ribosome collisions generated by YdiF(EQ_2_) are enriched for tmRNA. **(A)** MNase digestion followed by sucrose density gradient ultracentrifugation revealing increased presence of disome and trisome peaks indicative of ribosome collisions upon YdiF(EQ2) expression (top). Continuous histogram showing the average distribution of read lengths after size-selection and mapping. **(B)** Average RPF reads mapped to tmRNA (*ssrA*) in -YdiF(EQ_2_) and +YdiF(EQ_2_) cells. **(C)** Scatterplot showing the average TPM of RPFs ≥ 70 nt in -YdiF(EQ_2_) and +YdiF(EQ_2_) cells; perfect identity line is shown. **(D)** Bar graph showing the ribosome occupancy of tmRNA in -YdiF(EQ_2_) and +YdiF(EQ_2_) cells. Ribosome occupancy was calculated by dividing the normalized ribosome density of tmRNA on RPFs ≥ 70 nt in length in the ribo-seq library by the normalized expression of tmRNA in total RNA-seq. Error bars represent the SD of biological triplicate ribo-seq libraries. The *p-*value is the result of Welch’s two-tailed T-test.

Since collided ribosomes have a longer footprint than monosomes we next determined whether YdiF(EQ_2_)-expressing cells exhibit longer ribosome protected mRNA fragments. We grew cells to late logarithmic growth phase, snap-froze them in liquid nitrogen, and cryomilled with high magnesium to maintain the native positions of translating ribosomes (36). The lengths of the ribosome protected fragments (RPFs) recovered from both wild-type and YdiF(EQ_2_)-expressing cells centered around 25-40 nucleotides, corresponding to the approximate length of a monosome footprint and consistent with previous ribo-seq studies in bacteria (**Fig. 2A**) (36). Both strains also exhibited a second peak around 72-75 nucleotides, which is consistent with the length of a disome protected fragment. However, YdiF(EQ_2_)-expressing cells exhibited greater levels of longer ribosome protected fragments, further indicating that there are higher levels of ribosome collisions in these cells.

### Ribosome collisions generated by YdiF(EQ_2_) are enriched for tmRNA

Having determined that YdiF(EQ_2_) expression generates high levels of ribosome collisions (**Fig. 2A**), we next sequenced transcripts associated with collisions. We found that tmRNA (encoded by *ssrA*) was highly enriched in the ribosomes of YdiF(EQ_2_)-expressing cells compared to cells not expressing YdiF(EQ_2_) (**Fig. 2B, Fig. S1**). tmRNA was even more enriched in the ribosomes of YdiF(EQ_2_)-expressing cells when we compared only the longer (≥70 nucleotides) ribosome protected fragments that arise from ribosome collisions (**Fig. 2B**). Reads mapping to tmRNA were approximately 2-fold greater in ribosome collisions in YdiF(EQ_2_)-expressing cells (**Fig. 2C**). Strikingly, in these cells, tmRNA was among the top three most highly translated transcripts in all three biological replicates. The ribosome occupancy of tmRNA (which is a measure of ribosome density compared to cellular transcript level) in YdiF(EQ_2_)-expressing cells was significantly higher than in wild-type (3-fold higher ribosome occupancy, p = 0.028) (**Fig. 2D**) (44). These data indicate that tmRNA associates with ribosome collisions generated by YdiF(EQ_2_) expression.

### Ribosome collisions generated by YdiF(EQ_2_) are rescued by *trans-*translation

Reads mapping to tmRNA were highly enriched in ribosome collisions in YdiF(EQ_2_)-expressing cells, suggesting that tmRNA is recruited to collided ribosomes (**Fig. 2**). To further verify that ribosome collisions caused by YdiF(EQ_2_) generate substrates for *trans-*translation, we measured global *trans-*translation activity in cells expressing YdiF(EQ_2_). We replaced the degron-encoding open reading frame of tmRNA with a 6xHis tag so that tmRNA-mediated ribosome rescue will append a 6xHis tag to the nascent protein (**Fig. 3A**). We also mutated the C-terminal dialanine motif to a diaspartate motif so that the His-tagged nascent proteins are not targeted for degradation (24). We then harvested cell lysate from wild-type and YdiF(EQ_2_)-expressing cells and compared tagging levels by western blot using anti-His antibodies. As expected, even in wild-type cells, tmRNA tagging levels are high, which is consistent with previous measurements indicating that each ribosome participates in *trans*-translation two to three times per cell cycle (**Fig. 3A**, α-His western blot) (45,46). YdiF(EQ_2_) expression further increased levels of tmRNA-mediated tagging (**Fig. 3B**). Increased levels of tmRNA association with ribosome collisions coupled with the global increase in *trans*-translation activity in YdiF(EQ_2_) expressing cells further confirms that YdiF(EQ_2_) generates ribosome collisions that are rescued by *trans*-translation.

**Figure 3.**
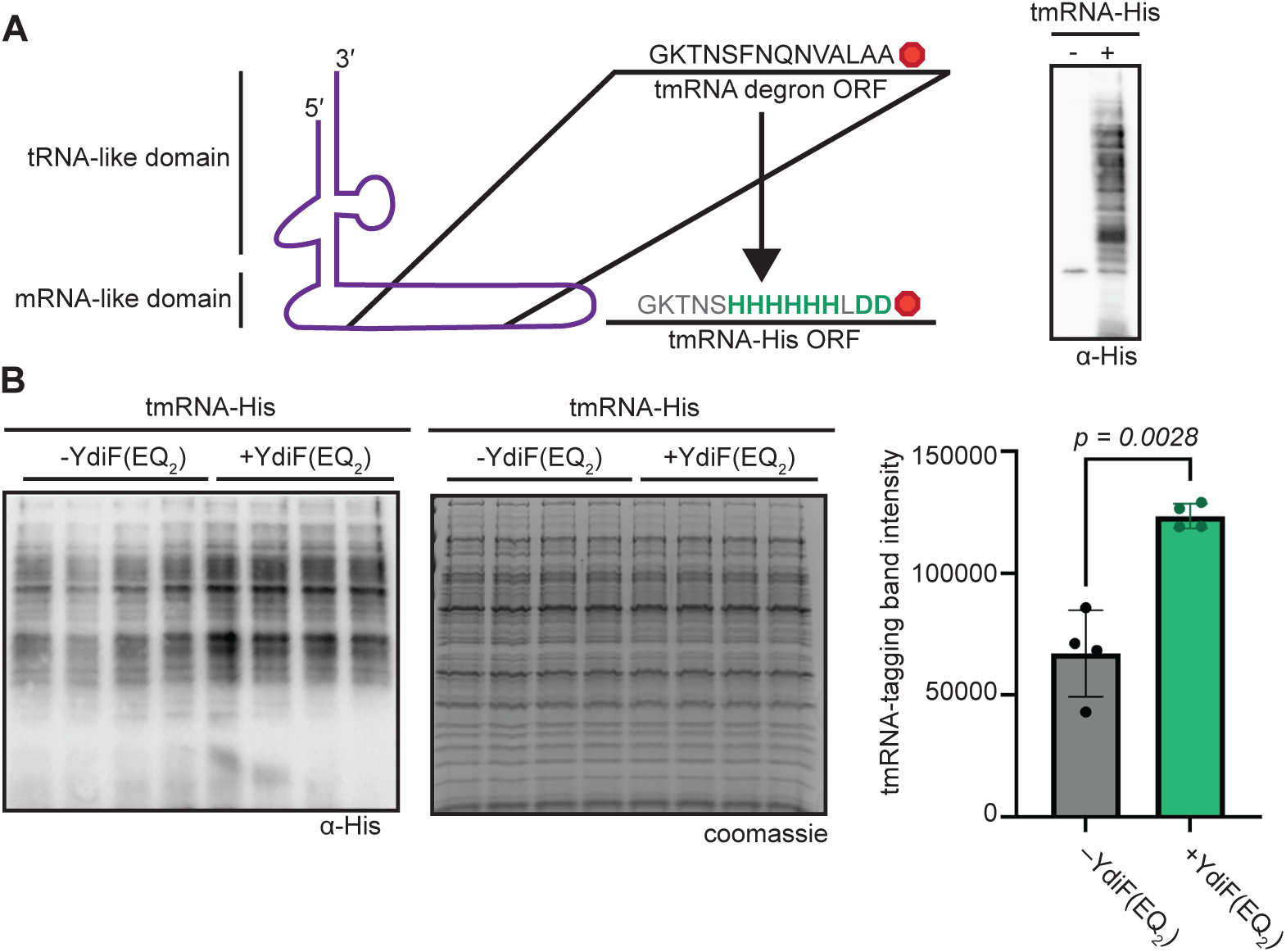
Ribosome collisions generated by YdiF(EQ_2_) are rescued by *trans-*translation. **(A)** Cartoon schematic illustrating how the degron tag of tmRNA was recoded to a 6x Histidine tag to monitor tmRNA-mediated tagging by western blot. Western blot of whole cell lysate from cells expressing only wild-type tmRNA or wild-type tmRNA and the recoded version. **(B)** Anti-His western blot showing tmRNA tagging levels in cells with and without YdiF(EQ_2_) expression. Four biological replicates are loaded. Coomassie gel shows total protein as a loading control. Quantification of tmRNA-tagging levels is shown in bar graph with error bars representing the standard deviation of biological quadruplicate. The *p*-value is the result of Welch’s two-tailed T-test.

### The Rae1 endonuclease generates substrates for *trans*-translation in response to ribosome collisions

tmRNA is recruited to YdiF(EQ_2_)-induced ribosome collisions (**Fig. 2**, **Fig. 3**). Since tmRNA only rescues ribosomes at the 3′ end of an mRNA, these data indicate that mRNA cleavage has occurred on these collided ribosomes (19,21,47–49). While *E. coli* uses SmrB to cleave mRNA on collided ribosomes, Gram-positive bacteria lack the SmrB nuclease (50). Therefore, we sought to determine the endonuclease responsible for increased *trans*-translation activity during YdiF(EQ_2_) expression. We hypothesized that the <u>R</u>ibosome <u>a</u>ssociated <u>e</u>ndonuclease Rae1 generates *trans*-translation substrates since numerous studies have shown that it cleaves mRNA on stalled ribosomes (51–54). To test this, we expressed YdiF(EQ_2_) in either wild-type cells or cells from which we deleted *rae1* and compared levels of tmRNA tagging. In the absence of Rae1, YdiF(EQ_2_)-expressing cells exhibit tmRNA tagging levels similar to the background levels exhibited by wild-type cells (**Fig. 4A**). These data indicate that Rae1 is the source of *trans*-translation substrates that are generated in response to YdiF(EQ_2_) expression. Altogether, these results support a model in which Rae1 is recruited to collided ribosomes and cleaves associated mRNA; Rae1-mediated truncation of mRNA then allows collided ribosomes to be rescued by *trans*-translation (**Fig. 4B**)

**Figure 4.**
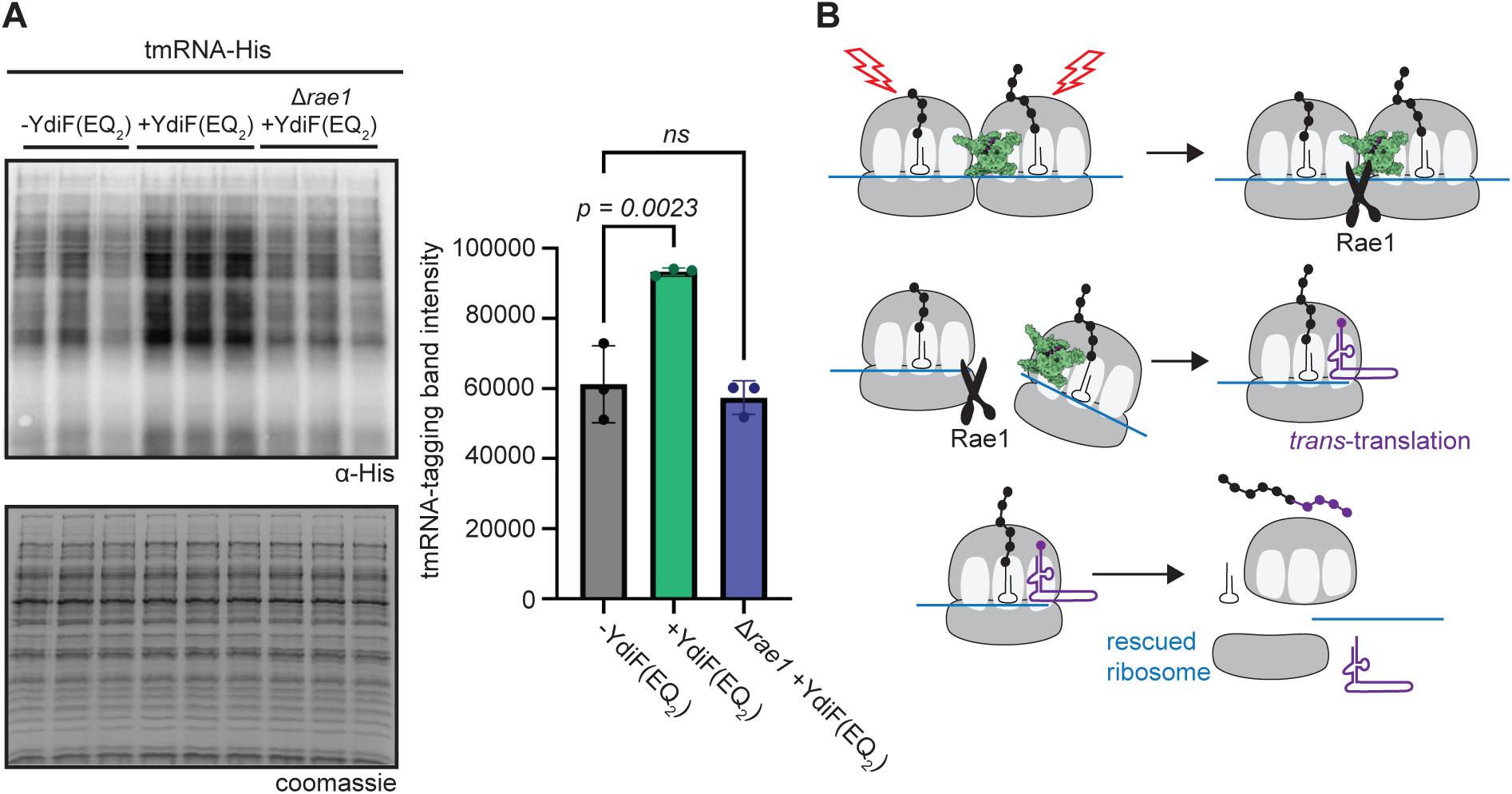
The Rae1 endonuclease generates substrates for *trans*-translation in response to ribosome collisions. **(A)** Coomassie gel, western blot, and western band intensity quantification analysis of *trans-*translation activity in -YdiF(EQ_2_), +YdiF(EQ_2_), and *Δrae1*+ YdiF(EQ_2_) cells. Error bars represent the standard deviation of biological triplicate. *p*-values are the result of one-way ANOVA and Dunnett’s multiple comparisons test. **(B)** Cartoon schematic showing proposed model for Rae1 and *trans*-translation mediated rescue.

The Rae1 endonuclease is required for increased tmRNA tagging during YdiF(EQ_2_)-induced ribosome collisions (**Fig. 4A**). To determine whether Rae1 contributes to the baseline level of *trans-*translation observed in wild-type cells without any translation stress, we compared tmRNA tagging levels in wild-type cells and Δ*rae1* cells. In the absence of translation stress, tmRNA tagging levels in wild-type cells and Δ*rae1* cells were not significantly different (**Fig. S2**). These data indicate that Rae1 does not contribute to the baseline activity of *trans*-translation in wild-type cells, pointing toward its specialization for cleaving mRNA on collided ribosomes.

### *trans-*Translation is important for fitness in response to ribosome collisions

YdiF(EQ_2_)-induced ribosome collisions significantly increase *trans*-translation activity (**Fig. 4**). Therefore, we next asked whether the genes encoding *trans*-translation components are important for fitness during YdiF(EQ_2_) expression. *trans*-Translation is mediated by tmRNA (encoded by *ssrA*) and its partner protein SmpB. As observed previously, YdiF(EQ_2_) expression has only a minor effect on growth on plates in the wild-type background (**Fig. 1**, **Fig 5A**). In contrast, Δ*ssrA-smpB* cells expressing YdiF(EQ_2_) exhibit a four-log decrease in growth on plates (**Fig. 5A**). Growth in LB liquid culture was also severely impaired (**Fig. 5B**). The log-phase growth rate of YdiF(EQ_2_)-expressing cells was significantly decreased in cells lacking *trans*-translation. Similarly, time to exit lag phase increased (**Fig. 5B**). These results indicate that *trans*-translation is important for fitness when cells are experiencing widespread ribosome collisions.

**Figure 5.**
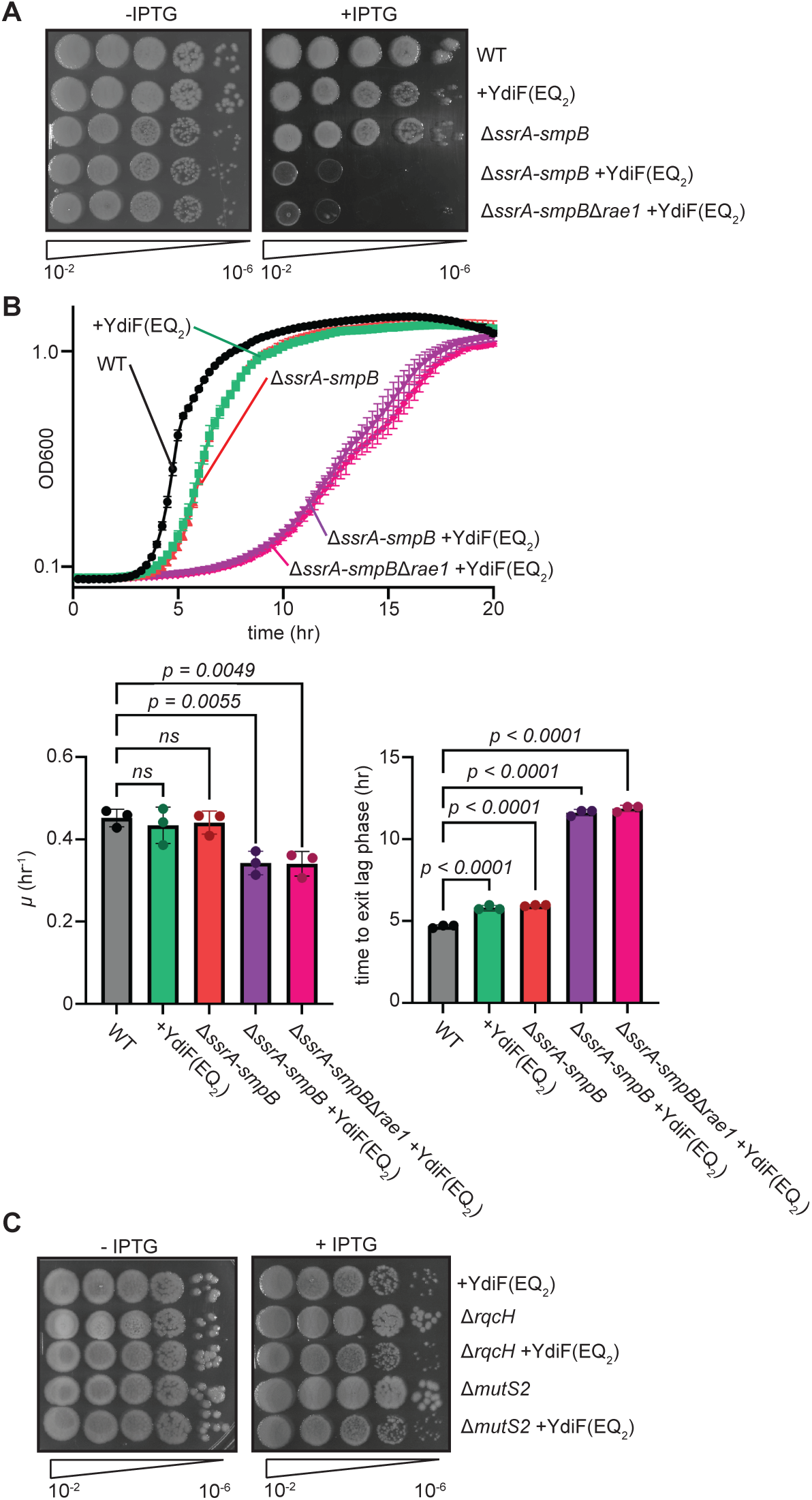
*trans*-translation is required to support growth when ribosome collisions are generated via YdiF(EQ_2_) expression. **(A)** Serial dilutions of cells expressing YdiF(EQ_2_) in combination with deletions of the genes encoding *trans-*translation and Rae1. **(B)** (top) Growth curves of cells expressing YdiF(EQ_2_) in combination with deletions of genes encoding *trans-*translation and Rae1. Error bars represent the SEM of biological triplicate growth curves. (Bottom left) Specific growth rate for growth curves. (Bottom right) Time to exit lag phase for growth curves (measured as time at which OD exceed 0.125 when starting from OD600=0.0005 diluted from overnight culture). Error bars represent the standard deviation of biological triplicate. *p*-values are the result of one-way ANOVA and Dunnett’s multiple comparisons test. **(D)** Serial dilution of cells expressing YdiF(EQ_2_) in combination with deletions of *rqcH* and *mutS2*. Experiment was performed in biological triplicate, and representative plate image is shown.

We observed that the Rae1 endonuclease is required to generate increased *trans*-translation activity upon YdiF(EQ_2_) overexpression (**Fig. 4A**). We therefore hypothesized that deletion of *rae1* would rescue the growth of Δ*ssrA-smpB* cells expressing YdiF(EQ_2_), reasoning that fewer *trans*-translation substrates would decrease the need for *trans*-translation. However, YdiF(EQ_2_) overexpression negatively impacted the growth of Δ*ssrA*-*smpB*Δ*rae1* cells to the same extent as Δ*ssrA*-*smpB* cells (**Fig. 5A, 5B**). Therefore, *trans*-translation is important for fitness during ribosome collisions, even when Rae1 is absent.

To determine whether other ribosome rescue factors are important for fitness during ribosome collisions generated by YdiF(EQ_2_), we overexpressed YdiF(EQ_2_) in Δ*rqcH* and Δ*mutS2* cells. Growth of both Δ*rqcH* and Δ*mutS2* cells expressing YdiF(EQ_2_) was comparable to the modest growth defect observed in wild-type cells expressing YdiF(EQ_2_) (**Fig. 5C**). Loss of RQC or a splitting factor did not exacerbate the growth defect induced by YdiF(EQ_2_) expression. Therefore, these data indicate that RqcH and MutS2 are not important for fitness during ribosome collisions generated by YdiF(EQ_2_).

### Expression of other housekeeping ABCF ATPase(EQ_2_) mutants also increases ***trans***-translation activity

*B. subtilis* encodes three additional housekeeping ABCF ATPases: YfmR, YfmM, and YkpA (29,32,33). To test whether increased *trans-*translation activity was specific to the expression of YdiF(EQ_2_) or is caused by expression of any ATPase defective ABCF ATPase, we used the tmRNA-His strain to examine *trans*-translation activity during expression of YfmR(EQ_2_), YfmM(EQ_2_), and YkpA(EQ_2_). All four ABCF(EQ_2_) mutants significantly stimulated *trans*-translation activity (**Fig. S3**). These results suggest that increased *trans*-translation is not specific to YdiF(EQ_2_) expression but is a general response to ribosome stalling.

### Erythromycin exposure increases *trans*-translation activity in a Rae1-dependent manner

We next determined whether Rae1 and *trans*-translation can be used to rescue antibiotic-induced ribosome collisions. Erythromycin binds 23S rRNA near the entrance to the exit tunnel and preferentially stalls the ribosome at R/K-X-R/K motifs (37,38,55). Upstream ribosomes that haven’t encountered an R/K-X-R/K motif will collide with these stalled ribosomes. Therefore, subinhibitory erythromycin treatment is a standard method used to induce ribosome collisions (13,18).

To test whether exposure to erythromycin increases *trans*-translation activity, we treated cells containing the tmRNA-His allele with a series of increasing erythromycin concentrations. We found that increasing erythromycin concentrations led to increased *trans*-translation activity in a titratable manner (**Fig. 6A**). To test whether increased *trans*-translation activity during exposure to erythromycin was dependent on Rae1, we deleted *rae1* in cells containing the tmRNA-His allele and treated them with 30 ng/µL of erythromycin. Whereas wild-type cells with the tmRNA-His allele exhibited increased *trans*-translation activity during exposure to erythromycin, *trans-*translation activity in Δ*rae1* cells did not increase in response erythromycin treatment (**Fig. 6B**). These data suggest that exposure to erythromycin also stimulates ribosome rescue through Rae1-mediated cleavage and *trans-*translation and demonstrate the general applicability of this pathway to rescue ribosome collisions.

**Figure 6.**
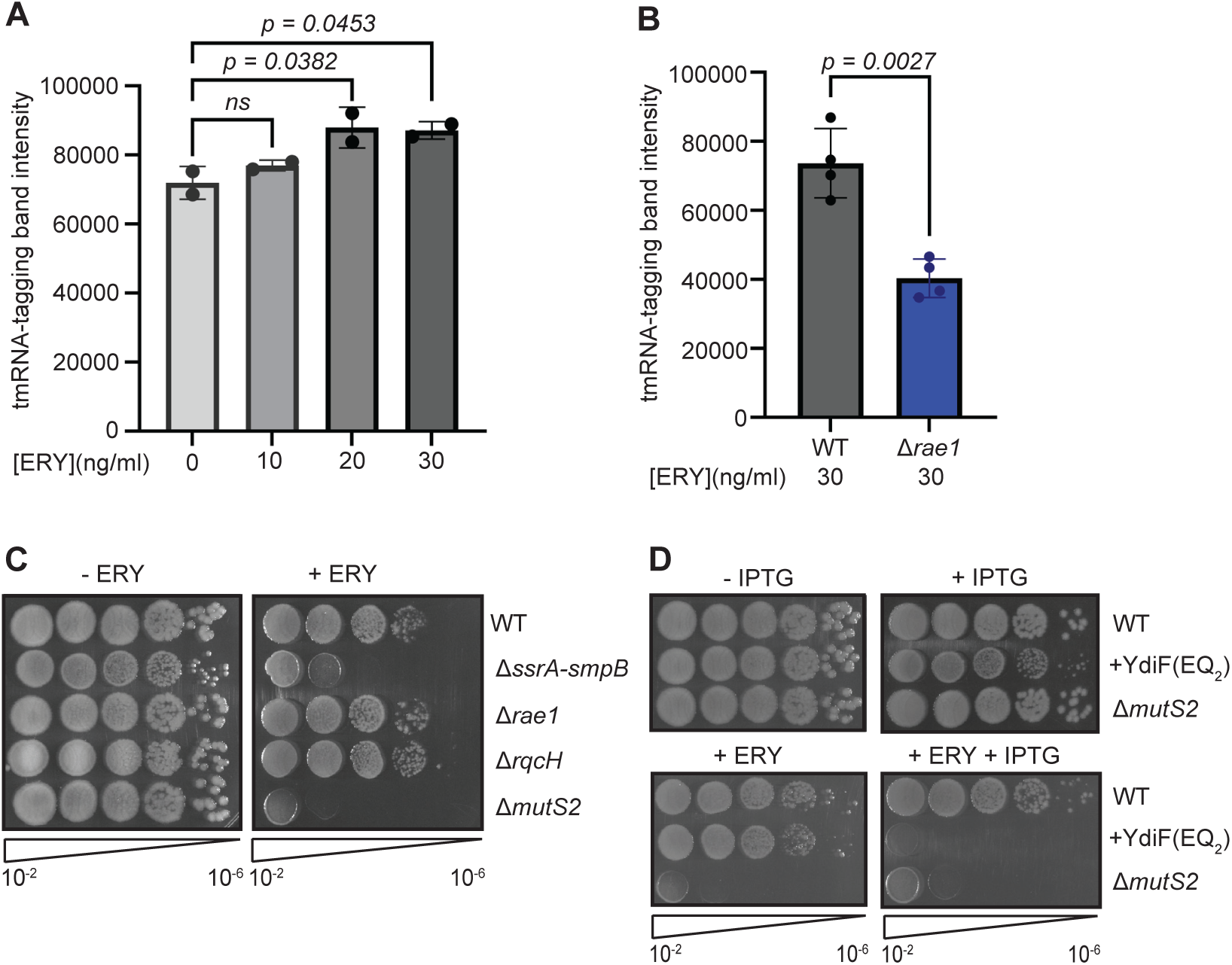
Erythromycin exposure increases *trans*-translation activity in a Rae1-dependent manner. **(A)** tmRNA-tagging activity in DT350 cells treated with increasing doses of erythromycin as determined by western blot. Error bars represent the standard deviation of biological duplicate. *p-*values are the result of one-way ANOVA and Dunnett’s multiple comparisons test. (**B)** tmRNA-tagging activity in wild-type and *Δrae1* cells treated with 30 ng/ml erythromycin. Error bars represent the standard deviation of biological quadruplicate. The *p-*value is the result of Welch’s two-tailed T-test. **(C)** Serial dilution of wild-type, *ΔssrA-smpB, Δrae1, ΔrqcH,* and *ΔmutS2* cells with and without 40 ng/ml erythromycin. **(D)** Serial dilution of wild-type, +YdiF(EQ_2_), and *ΔmutS2* cells on plates containing 40 ng/ml erythromycin. Experiment was performed in biological triplicate, and representative plate image is shown.

Since erythromycin-induced ribosome collisions invoke tmRNA/Rae1-mediated rescue, we compared erythromycin sensitivity of YdiF(EQ_2_)-expressing cells to cells harboring deletions of ribosome rescue pathways that respond to ribosome collisions. Consistent with previous work, cells lacking *trans*-translation or the ribosome splitter MutS2 were more sensitive to erythromycin-induced collisions (**Fig. 6C**) (13,14,56,57). Cells expressing YdiF(EQ_2_) were also highly sensitized to erythromycin, exhibiting a fitness defect that was even more severe than Δ*mutS2* cells (**Fig. 6D**). These data suggest that the ribosome stalling and collisions induced by erythromycin treatment and YdiF(EQ_2_) expression are additive, leading to compounding fitness defects.

### Rae1 is distributed among diverse bacterial phyla

Rae1 and *trans*-translation respond to ribosome collisions that result from diverse causes (**Fig.4**, **Fig. 6**). To investigate the conservation and prevalence of the components of this pathway outside of *B. subtilis*, we surveyed 18,663 representative reference genomes to determine the distribution of collision rescue pathways. The genes encoding *trans*-translation (*ssrA* and *smpB*) are conserved in >97% of surveyed bacterial genomes, consistent with the fact that non-stop mRNA is a major problem in all bacteria (**Fig. 7A**) (50,58,59). Both the collision rescue pathway described here, and the analogous rescue mechanism recently described in *E. coli* converge on *trans*-translation for ribosome rescue (18). Therefore, the necessity of *trans*-translation can be viewed not only as a non-stop rescue mechanism, but also as a ribosome collision rescue mechanism in diverse bacteria. The genes encoding components of the RQC pathway (*mutS2* and *rqcH*) are broadly distributed throughout the bacterial domain but are not as universally conserved as *trans*-translation (**Fig. 7A**).

**Figure 7.**
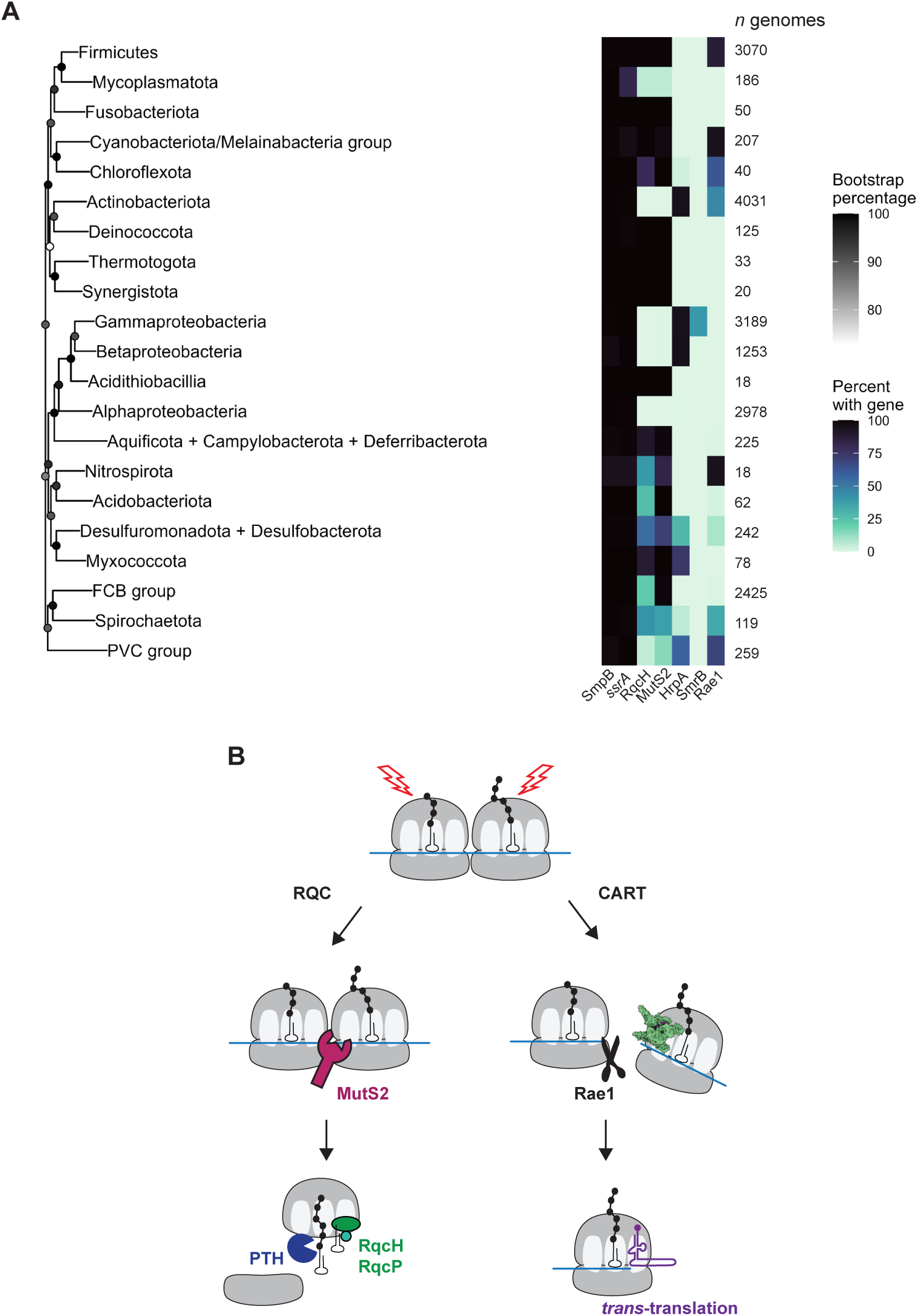
Distribution of genes encoding the RQC and CART pathways. **(A)** Conservation of ribosome rescue pathways across >18,000 bacterial genomes. Heatmap showing the percentage of genomes per phyla containing genes encoding SmpB, *ssrA*, RqcH, MutS2, HrpA, SmrB, and Rae1. NCBI accession numbers of all surveyed genomes and the query sequences used to search for each gene are given in the GitHub repository in the Data Availability Statement. Bootstrap percentages are shown as darkened circles at each node. **(B)** Schematic overview of RQC vs. CART pathways for rescuing collided ribosomes.

Rae1 is broadly distributed throughout the bacterial domain and is extremely well conserved in Firmicutes, Cyanobacteria, and Nitrospirota; Rae1 is found in greater than 87%, 95%, and 94% of surveyed genomes in these phyla, respectively (**Fig. 7A**). We additionally detected intermediary Rae1 conservation in Actinobacteria (45%), Chloroflexota (63%), and Spirochaetota (34%). SmrB is an unrelated but analogous nuclease that associates with collided ribosomes and generates *trans*-translation substrates in *E. coli* (18). SmrB exhibits a much more limited distribution than Rae1 and is restricted to less than half of the surveyed Gammaproteobacteria (39%) and a handful of Betaproteobacteria (**Fig. 7A**). Although Rae1-mediated mRNA cleavage has so far only been reported in *B. subtilis*, its conservation suggests that many additional phyla may use Rae1 as a strategy to make collided ribosomes available for rescue by *trans-*translation.

In addition to SmrB/tmRNA-mediated collision rescue, *E. coli* also uses the RNA helicase HrpA to rescue stalled ribosomes (16,60). HrpA exerts a pulling force on the mRNA, causing dissociation of the subunits of the stalled ribosome. Although the fate of the resultant obstructed 50S subunit remains unclear, recent work suggests that the obstructed 50S rejoins a 30S subunit and that the resulting 70S can be rescued by *trans*-translation (61). HrpA is well conserved in surveyed Beta- and Gammaproteobacteria (96% and 95% respectively), as well as in Actinobacteria (95%) (**Fig. 7A**). HrpA is also present in surveyed Desulfuromonadota (28%) and Myxococcota (74%) genomes. Interestingly, we observed limited overlap in the conservation of Rae1 and HrpA (except in Actinobacteria). While most phlya encode either SmrB, Rae1, or HrpA, the absence of any homolog of these proteins in remaining phyla raises the exciting possibility that these bacteria use another nuclease or another strategy entirely to incorporate *trans*-translation into ribosome collision rescue.

## Discussion

Our data support a model in which Rae1 cleaves mRNA on collided ribosomes, making these ribosomes available for rescue by *trans*-translation. We find that tmRNA associates with collided ribosomes and that tmRNA-mediated tagging increases globally in response to widespread collisions in *B. subtilis* (**Fig. 2**, **Fig. 3**). Increased tmRNA tagging activity is dependent on the ribosome associated endonuclease Rae1 (**Fig. 4**), indicating that Rae1 is responsible for generating the truncated mRNA that is required for ribosome targeting to *trans*-translation. We detected Rae1-dependent tmRNA-tagging in response to ribosome collisions induced by overexpression of an inactivated ribosome binding protein and during erythromycin treatment (**Fig. 6**), suggesting that this mechanism may respond to ribosome collisions that arise from diverse causes. In communication with the Braun and Condon labs, we refer to this mechanism as <u>C</u>ollision-<u>A</u>ssociated <u>R</u>ae1-induced *trans-*Translation (CART).

How does Rae1 recognize stalled and collided ribosomes? Structures of ribosome collisions from *E. coli*, *B. subtilis*, yeast, and humans reveal a similar interaction interface, with the majority of contacts occurring between the small subunits of the collided ribosomes (14,62). Therefore, collided ribosomes do not form random contacts but produce a platform that has a conserved architecture. Collision rescue factors likely evolved to specifically recognize the collision interface, since they often interact with the same small subunit proteins. For example, the SMR domain of the collision rescue factors MutS2 and SmrB both interact with uS7 and uS11 on the stalled ribosome and with uS3 on the collided ribosome (14,18). Rae1 does not possess an SMR domain, and there are presently no structures of ribosome-bound Rae1 since its association with ribosomes is transient and therefore challenging to capture (51). Elegant work by Deves and colleagues determined that Rae1 cleaves mRNA approximately 12 nucleotides upstream of the A site codon in a stalled ribosome, indicating that, like SmrB, Rae1 cleaves mRNA upstream of the stalled ribosome (52). Indeed, Deves *et al.* noted that the 12-nucleotide spacing between the A site and the mRNA cleavage site is consistent with Rae1 contacting a trailing ribosome and therefore proposed that Rae1 participates in collision rescue (52). In the future, structural characterization of Rae1 binding to collided ribosomes is necessary to determine its precise mode of recognition.

If Rae1 cleaves mRNA between the stalled and collided ribosome, what is the fate of the stalled ribosome? Loss of ribosome occupancy on the mRNA downstream from the collision site leads to deprotection and degradation of the mRNA, since translational efficiency of an mRNA affects its stability (63–67). Without ribosome protection of the mRNA downstream of the stalled ribosome, degradation factors have increased access to the mRNA (68–70). Therefore, one possibility is that the mRNA fragment that arises post-Rae1 cleavage is degraded by exonuclease activity until no mRNA remains in the A site of the stalled ribosome, allowing it to be rescued by *trans*-translation. Therefore, *trans*-translation is likely able to rescue both the stalled and collided ribosome following Rae1 cleavage. Experiments that individually track the stalled ribosome will be critical for determining the mode of its rescue.

Ribosome collisions in *B. subtilis* are also rescued by the splitting factor MutS2 (RqcU). MutS2 specifically recognizes collided ribosomes and uses its ATPase activity to pry apart the large and small subunit of the stalled ribosome (13,14). Splitting of the subunits leaves the large 50S subunit obstructed with peptidyl-tRNA. Template-free addition of an alanine tag to the nascent chain by RqcH and its processivity proteins RqcP/YlmH allows the peptidyl tRNA to slip out of the exit tunnel, where the cytoplasm-exposed ester linkage is then cleaved by PTH (17). Although MutS2 encodes an SMR domain that it uses to interact with the disome interface, it lacks key residues that are required for nuclease activity (14). Therefore, unlike SmrB or Rae1, MutS2 does not generate truncated mRNA on collided ribosomes and therefore does not trigger subsequent rescue by tmRNA. Consistent with previous work, we find that MutS2 and Rae1 act in independent pathways and likely have independent targets (52). Recent work by the Braun and Condon labs revealed a subset of mRNAs that are known to stall the ribosome (such as *spyA* and *bmrX*) and are cleaved by Rae1 (52,54,71). Rates of cleavage of these mRNAs is unaffected by loss of MutS2 (71), indicating that ribosomes stalled on these transcripts are targeted by Rae1, but not MutS2. Similarly, we observed that while YdiF(EQ_2_)-expressing cells require tmRNA for fitness, they do not require MutS2 (**Fig. 5**), suggesting that MutS2 does not play a major role during YdiF(EQ_2_)-induced ribosome stalling. Since the RQC pathway works independently of Rae1 or *trans*-translation, we propose to use <u>C</u>ollision-<u>A</u>ssociated <u>R</u>ae1-induced *trans-*Translation (CART) to describe the truncation and rescue pathway mediated by Rae1 and tmRNA. Future studies will investigate which types of ribosome stalling leads to rescue by RQC vs. CART.

The SmrB endonuclease found in Gammaproteobacteria participates in a rescue pathway that is analogous to what we describe here for Rae1 (18). However, Rae1 and SmrB are unrelated nucleases, and they differ in their phylogenetic distribution (**Fig. 7**). SmrB is restricted mainly to Gammaproteobacteria, whereas Rae1 is much more broadly distributed (**Fig. 7**). Interestingly, SmrB is found in bacteria that lack the RQC pathway, whereas Rae1 is found in phyla that use RQC for ribosome rescue. Why maintain Rae1 in the presence of RQC? One possibility is that destruction of ribosome-stalling mRNA is advantageous. Indeed, this has been shown for the Rae1-targeted *spyT* transcript, which encodes a toxic peptide (54). It is easy to imagine additional scenarios where mRNA degradation could be more beneficial than ribosome splitting. For example, infection with a phage that differs in codon usage could result in ribosome stalling and collisions when phage-encoded transcripts are translated. In such a scenario, Rae1 or SmrB could defend against mRNA from an invader. The importance of degrading mRNAs on which ribosomes become stalled is further highlighted by the no-go decay pathway in eukaryotic ribosome quality control (72,73). In no-go decay, mRNA on stalled ribosomes is first cleaved by an endonuclease, then followed by exonuclease digestion of the mRNA. In yeast, no-go decay can be initiated by the endonuclease Cue2, which cleaves mRNA on collided ribosomes in a manner similar to what we propose for Rae1 (74–76).

The importance of *trans*-translation in ribosome rescue is underscored by the fact that genes encoding *trans*-translation are conserved in nearly all bacterial species (**Fig. 7**) (50,58,77). It is striking that numerous pathways for ribosome rescue collisions converge on *trans*-translation, and these pathways are not limited to nucleases like SmrB and Rae1. For example, the RNA helicase HrpA was recently proposed to split stalled ribosomes in such a way that 70S monosomes can reform in the absence of mRNA and be rescued by tmRNA (61). Collisions between ribosomes and stalled RNA polymerase also generate truncated mRNA, which the ribosome translates to the 3′ end before being rescued by *trans*-translation (78). Altogether, these findings demonstrate how diverse mechanisms that respond to problems with transcription and translation converge on *trans*-translation, further supporting its central role in ribosome quality control in bacteria.

## Materials and Methods

### Strains and media

The wild-type strain background is *B. subtilis* 168 *trpC2* (strain DT01) (79). All experiments were performed in LB media at 37°C with the addition of IPTG or erythromycin as specified.

Expression of YdiF(EQ_2_) was accomplished by integrating a plasmid, pDR111, containing the mutant YdiF(EQ_2_)-3xFLAG fused to an IPTG-inducible promoter P_hyper-spank_ and a weak RBS to minimize uninduced expression at the *amyE* locus, producing strain DT195. The glutamate residues to mutate were determined by multiple sequence alignment of all housekeeping ABCF ATPases from *E. coli* and *B. subtilis* and identification of their highly conserved ATP-binding cassettes (39). The same process was used to generate strains expressing YdiF-3xFLAG and YfmR-3xFLAG, producing strains DT194 and DT277 respectively. All plasmids were checked via whole plasmid sequencing by Plasmidsaurus using Oxford Nanopore Technology (Oxford Nanopore, R10.4.1). For all pDR111 transformants, stable integration was confirmed by starch hydrolysis test and α-FLAG western analysis for the expression product.

The tmRNA-His strain was generated by integrating recoded *ssrA* at the *sacA* locus on the chromosome. The 14-codon degron ORF was recoded to a hexahistidine tag producing DT350 (80). tmRNA-His strains expressing YdiF(EQ_2_)-3xFLAG, YfmR(EQ_2_)-3xFLAG, YfmM(EQ_2_)-3xFLAG, and YkpA(EQ_2_)-3xFLAG were generated by integrating pDR111 plasmid vectors into the *amyE* locus of the tmRNA-His strain as described above, producing strains DT354, DT365, DT367, and DT369 respectively.

Strain DT356, Δ*rae1::kan^R^* tmRNA-His + YdiF(EQ_2_)-3xFLAG, was obtained by first transforming genomic DNA from Δ*rae1::kan^R^* cells in the BKK collection (81) into the lab background, producing strain DT388. Next, the YdiF(EQ_2_)-3xFLAG expression plasmid (pDR111) was integrated into the *amyE* locus of Δ*rae1::kan^R^* cells. Finally, the tmRNA-His plasmid (ECE174) was integrated into the *sacA* locus of the Δ*rae1::kan^R^* + YdiF(EQ_2_)-3xFLAG cells. The tmRNA-His plasmid (ECE174) was also integrated into the *sacA* locus of Δ*rae1::kan^R^* cells, yielding DT375.

Similarly, strains DT180 and DT383 were obtained by first transforming genomic DNA from Δ*mutS2::kan^R^* and Δ*rqcH::kan^R^* cells in the BKK collection, respectively, into the lab background. Next, the YdiF(EQ_2_)-3xFLAG expression plasmid (pDR111) was integrated into the *amyE* locus of DT180 and DT383 cells, producing strains DT386 and DT384 respectively. DT395 was produced by first transforming markerless Δ*mutS2* cells with genomic DNA from Δ*rae1::kan^R^* cells in the BKK collection, producing Δ*mutS2*Δ*rae1::kan^R^*cells. Next, the YdiF(EQ_2_)-3xFLAG expression plasmid (pDR111) was integrated into their *amyE* locus, finally yielding DT395.

The Δ*ssrA-smpB::tet^R^* mutant was constructed by integrating a plasmid (KC249, Twist, digested with PvuI-HF) carrying the Δ*ssrA-smpB::tet^R^* deletion into the lab background, yielding strain DT364. The Δ*rae1::kan^R^* deletion was added to the Δ*ssrA-smpB::tet^R^* background by transforming genomic DNA from Δ*rae1::kan^R^* cells in the BKK collection, yielding strain DT371, and expression of YdiF(EQ_2_)-3xFLAG in these backgrounds was accomplished by integrating the YdiF(EQ_2_)-3xFLAG expression plasmid (pDR111) into the *amyE* locus of Δ*ssrA-smpB::tet^R^* and Δ*ssrA-smpB::tet^R^* Δ*rae1::kan^R^* cells, producing strains DT361 and DT373 respectively.

### Growth curves

Cells from fresh colonies were grown overnight in LB at 30°C then back-diluted to OD600=0.0005 in LB containing 2mM IPTG. The diluted cells were grown for 24 hours at 37 °C shaking constantly at 2 mm amplitude in Thermo Scientific 96-well flat-bottom plates (cat. no. 167008). OD600 values were measured every 15 minutes using a BioTek Synergy H1 microplate reader, Gen5 3.11. Growth curves were measured in biological and technical triplicate and error bars represent the SEM. Specific growth rate was calculated for each biological replicate using two OD600 values 1 hour apart in exponential growth phase via the formula *µ*=(ln(OD_2_) – ln(OD_1_))/(t_2_-t_1_) and statistical analysis was conducted using a one-way ANOVA and Dunnett’s multiple comparisons post-hoc test in GraphPad Prism v11.0.0.

### SDS-PAGE and western analysis

Cells were harvested via 1 minute centrifugation at 8,600 RPM. The supernatant was immediately pipetted off, and the pellet was stored at −80°C. The pellets were later thawed, resuspended in 100 μL of a solution of 10mM TE buffer (10mM Tris pH 8.0, 1mM EDTA) with 50mM EDTA and 1 μg/mL lysozyme, and incubated for 5 minutes at 37°C. 200 μL of 4X SDS loading buffer (6 mL Tris pH 6.8, 12 mL 220% SDS, 24 mL 50% glycerol, 800 μL BME, a pinch of bromophenol blue) were added to the lysed pellets, which were then boiled at 90°C for 4 minutes and placed on ice after boiling. The prepared lysates were loaded onto a 12% polyacrylamide gel for electrophoresis. Gels were cut according to the length and width of the loaded lanes, then transferred to a PVDF membrane (BioRad Immun-Blot PVDF Membrane) of the same size. α-FLAG membranes were blocked in 3% bovine serum albumin in PBS-T overnight at 4°C, while α-His membranes were blocked at room temperature for 45 minutes. α-FLAG membranes were incubated in 1:10,000 α-FLAG-HRP (Sigma, cat. no. A8592) in blocking buffer for 2 hours at room temperature before washing with PBS-T, exposure to chemiluminescent reagent, and imaging on ChemiDoc MP (BioRad), while α-His membranes were incubated in 1:10,000 α-His-HRP (Invitrogen, cat. no. ma1-21315-hrp) in blocking buffer overnight at 4°C before washing with PBS-T and imaging. α-EF-Tu membranes were blocked for 45 minutes at room temperature, incubated in 1:10,000 polyclonal α-EF-Tu in blocking buffer overnight at 4°C, washed three times with PBS-T, then incubated in 1:10,000 α-rabbit-HRP (EMD Millipore, cat. no. 12-348) in PBS-T before washing and imaging.

### Sucrose density gradient ultracentrifugation

Cells were grown to OD600=1.0 in 30 mL of LB in 250 mL baffled flasks at 37°C before harvest by centrifugation at 10,000 RPM for 10 minutes (Beckman Coulter Avanti J-15R, rotor JA-10.100). Supernatants were decanted and pellets were stored immediately at −80°C. Pellets were later washed and resuspended with ice-cold gradient buffer (20 mM Tris, 60 mM NH_4_Cl_2_, 7.5 mM MgAoc, 6 mM 2-mercaptoethanol, 0.5 mM EDTA) and lysed by bead beating (Benchmark Scientific BeadBug 6) on maximum speed in 30 second cycles with 3 minutes on ice in between each cycle.

Sucrose density gradients were prepared using 10% and 50% sucrose dissolved in gradient buffer via a Biocomp Gradient Station (Biocomp Instruments). Ribosome concentrations in the purified lysates were approximated by A_260nm_ measurements (NanoDrop) and 200 μg of RNA was loaded onto each gradient. Gradients were centrifuged for 3 hours at 30,000 RPM in a SW-41Ti rotor. Gradients were harvested using a Biocomp Gradient Station, which also read their spectra during fraction collection. Gradient fractions for western analysis were precipitated overnight at −20°C in a mixture of 500 μL fraction and 1.5 mL 99.5% ethanol. The precipitated fractions were pelleted by centrifugation at 14,800 RPM at 4°C for 30 minutes, then the supernatants were discarded and the pellets were prepared as for lysates for SDS-PAGE.

### Bio-orthoganal noncanonical amino acid tagging (BONCAT)

Cells from fresh colonies were grown overnight in LB at 30°C then back-diluted to OD600=0.0005 in LB containing 2mM IPTG. The diluted cells were grown for 24 hours at 37 °C shaking constantly at 2 mm amplitude in Thermo Scientific 96-well flat-bottom plates (cat. no. 167008). OD600 values were measured every 15 minutes using a BioTek Synergy H1 microplate reader, Gen5 3.11. A subsection of cells was rapidly collected in biological triplicate from mid-log phase before returning the plate to the microplate reader.

The collected cells were pooled then transferred to pre-heated test tubes containing *L*-homoproparyglglycine (Vector Laboratories) at a final concentration of 500 μM to measure total protein synthesis. Cells were grown to incorporate HPG into nascent proteins for 15 minutes before pelleting, after which they were fixed in 4% formaldehyde in PBS for 15 minutes. The fixed cells were thoroughly washed with 3% BSA in PBS before resuspension and incubation in 0.5% Triton X-100 in PBS for 15 minutes for permeabilization. Finally, click reactions in the fixed and permeabilized cells were performed using AlexaFluor 488 and “Click-iT Cocktail” according to manufacturer protocols (Invitrogen). The clicked cells were mounted on 1% agarose in PBS pads and visualized via fluorescence microscopy on the Nikon ECLIPSE Ni-E microscope with GFP-FITC filter cube, with a 20 ms exposure time for GFP. Phase contrast and fluorescence channels were combined, background-subtracted, and quantified using ImageJ v1.54g.

### Disome and trisome profiling

Cells from fresh colonies were grown in LB at 37°C then back-diluted to OD600=0.05 in 30mL LB containing 2mM IPTG in 250mL baffled flasks. Flasks were shaken at 220 RPM at 37°C until cells reached OD600=1.5, at which point the entire volume was flash frozen in liquid nitrogen, broken into small pieces with a hammer, and transferred to a 50 mL conical tube. Each 50 mL frozen sample was split into approximately equal volumes, weighed, and mixed with 11% of the sample mass of 10x lysis buffer (200 mM Tris pH 8, 1M NH_4_Cl, 50 mM CaCl_2_, 1%NP-40, 4% Triton-X-100, 1.6 M MgCl_2_). The samples were then cryomilled for four cycles of 3 minutes at 15 hz. The samples were rechilled in liquid nitrogen between each cryomilling to ensure the samples remained frozen. A small quantity of each sample was set aside for RNA-seq. Once pulverized, the lysates were thawed and cleared by centrifugation at 10,000 RPM at 4°C for 10 minutes, then loaded onto a sucrose cushion (1.1 M sucrose, 20 mM Tris pH 8, 500 mM NH_4_Cl, 10 mM MgCl_2_, 0.5 mM EDTA, 6 mM BME). The cushioned lysates were then centrifuged at 60,000 RPM at 4°C for 2.5 hours in a TI-70 fixed angle rotor (Beckman Coulter Avanti J-15R). The supernatant was carefully decanted, then the pellet was washed and resuspended in resuspension buffer (20 mM Tris pH 8, 15 mM MgCl_2_, 100 mM NH_4_Cl, 5 mM CaCl_2_). The resuspended pellet was allowed to solubilize at room temperature on a roller drum for 5 minutes, at which point the A_260_ measurements of each sample were taken (NanoDrop) and micrococcal nuclease (New England Biolabs, cat. no. M0247S) equal to 2*((A_260_*30)/2000) μL was added to each along with 6 μL of Superase·In RNAse inhibitor (Thermo Fisher, cat. no. AM2696). The samples were then returned to the roller drum at room temperature for 45 minutes, after which they were centrifuged for 10 minutes at 20,000 RCF at 4°C and loaded onto 10%-50% sucrose gradients. The samples were finally centrifuged at 35,000 RPM at 4°C for 2.5 hours, after which the gradients were collected as described above and the monosome peaks were set aside and frozen at −80°C.

### Ribo-seq

Monosome RNA was extracted via phenol:chloroform:isoamyl alcohol precipitation (Thermo, cat. no. 15593031) and precipitated overnight at −80°C in an equal volume of isopropanol with 3 μL of glycoblue (Thermo, cat. no. AM9515). Ribosome protected fragments (RPFs) from monosomes were isolated via electrophoresis in a 15% TBE-urea 1.00 mm precast Criterion Gel (BioRad). One electrophoresed, the gel was shaken in 50 mL of 1x TBE running buffer with 10 μL sybergold for 10 minutes before the gel was imaged on a Typhoon FLA 7000 (General Electric Company) to guide cutting of RNA between approximately 15 nt and 60 nt. The gel slices were extruded via centrifugation at max speed, then resuspended in 500 μL RNA elution buffer (300 mM NaOAc pH 5.5, 1 mM EDTA pH 8) and 2 μL Superase·In RNAse inhibitor before rotation overnight at 4°C. Finally, the RPFs were then precipitated with isopropanol at −80°C before final resuspension in 15 μL 1x PNK buffer (New England Biolabs, cat. no. M0201) with 0.5 μL Superase·In RNAse inhibitor.

RPFs libraries were prepared using a NEBNext Small RNA library kit and sequenced on a NextSeq500 HO-150 flowcell (Illumina) with 2×75bp paired-end reads at the Cornell Transcriptional Regulation and Expression (TREx) Facility. Reads were trimmed in Geneious Prime v.2025 2.2 using the NEBNext Small RNA library kit adaptors as guides. Reads less than 23 nt in length were discarded (36). Reads were then assembled to the *B. subtilis* 168 reference genome from NCBI (GCF_000009045.1), sample TPMs were calculated, and mapped read lengths and abundances were quantified. Sample TPMs were compared in RStudio v.2025.09.1+401 using R v.4.5.1. All code used for Ribo-seq and translational efficiency analysis is available on https://github.com/danieltetreault7289/CART-riboseq-phylogenetics.

### tmRNA-His tagging to measure *trans-*translation activity

Cells from fresh colonies were grown overnight in LB at 30°C then back-diluted to OD600=0.05 in 30mL LB containing 2mM IPTG in 250mL baffled flasks. Flasks were shaken at 220 RPM at 37°C until cells reached OD600=1.0, at which point 1mL of cells was harvested for SDS-PAGE. Cell harvests were volume-adjusted to be OD600-equivalent with the first culture harvested. Cells were lysed and prepared for SDS-PAGE as described above. Each lysate was run on a 12% SDS-PAGE gel, Coomassie stained, and quantified for total protein content by band intensity quantification in ImageJ v1.54g to normalize loading volumes to the densest sample. Then, another SDS-PAGE gel of each lysate was run with normalized loading volumes, transferred to a PVDF membrane, and blotted as described above. tmRNA-His-tagging band intensity quantification was performed in ImageJ v1.54g.

### Gene detection

Genomes and annotated protein sequences from bacterial reference genomes (18,663 genomes) were downloaded from the NCBI RefSeq database on 30 April 2026. In the NCBI RefSeq database, reference genomes serve as a representative for the bacterial species, and approximately one reference genome is assigned to each species. Species with reference genomes that had >80% CheckM completeness and <10% CheckM contamination scores were included in our dataset (82). Protein sequences were detected using HMMER v3.3 (hmmsearch) (hmmer.org) with an E-value cutoff of 1×10^-10^. Query sequences used to build profiles for each protein are available on https://github.com/danieltetreault7289/CART-riboseq-phylogenetics. When querying for HrpA, both annotated HrpA and HrpB proteins were detected. To filter out HrpB, we applied a size filter to select only detected proteins >1200 amino acids in length. Methods for the nHMMER search for *ssrA* and descriptions of filtering criteria for RqcH, MutS2, and SmrB can be found in (50). All R code used to detect ribosome rescue factor genes is available at https://github.com/danieltetreault7289/CART-riboseq-phylogenetics.

### Phylogenetics

A maximum-likelihood tree of one random genome in each of the 21 bacterial phyla was built from SNPs in the universal gene set, which contains 16 universally conserved genes, using gToTree v1.8.17 (83). Taxonomic classification was assigned to the genomes using the NCBI Taxonomy database (84) and taxonkit v0.17.0 (85). Phyla were named using the conventions in Coleman *et al.* (86). The tree was visualized using ggtree v.4.3.0 (87). All R code used to build and visualize the tree is available at https://github.com/danieltetreault7289/CART-riboseq-phylogenetics.

## Acknowledgements

H.A.F., D.D.T., C.R.P., and K.C. were supported by NIH R35GM147049. D.D.T. and C.R.P. were supported by Graduate Research Fellowships from the National Science Foundation. We would like to thank Kevin England for assistance with BONCAT and the markerless Δ*mutS2* strain. We thank the Cornell TREx facility for sequencing the RPF libraries. We thank Gisela Storz, Allen Buskirk, Frédérique Braun, Ciaran Condon, Gene-Wei Li, and Kelley Gallagher for helpful discussions.

## Data Availability

Ribo-seq and RNA-seq data are available on the NCBI Gene Expression Omnibus (GEO) under accession numbers GSE342018 and GSE342019 respectively. csv files including global translatome RPFs (reported in TPM) and RPFs ≥70 nt, as well as ribosome occupancy scores for RPFs ≥70 nt can be found on GEO (GSE342018). Genome accession numbers and presence or absence of each rescue factor, gene sequences used to build HMMER profiles, random genomes in each of the 21 bacterial phyla used to build a maximum-likelihood tree, and all R code used to analyze Ribo-seq data, detect ribosome rescue factor genes, and build the phylogeny are available at https://github.com/danieltetreault7289/CART-riboseq-phylogenetics.

**Table 1.**
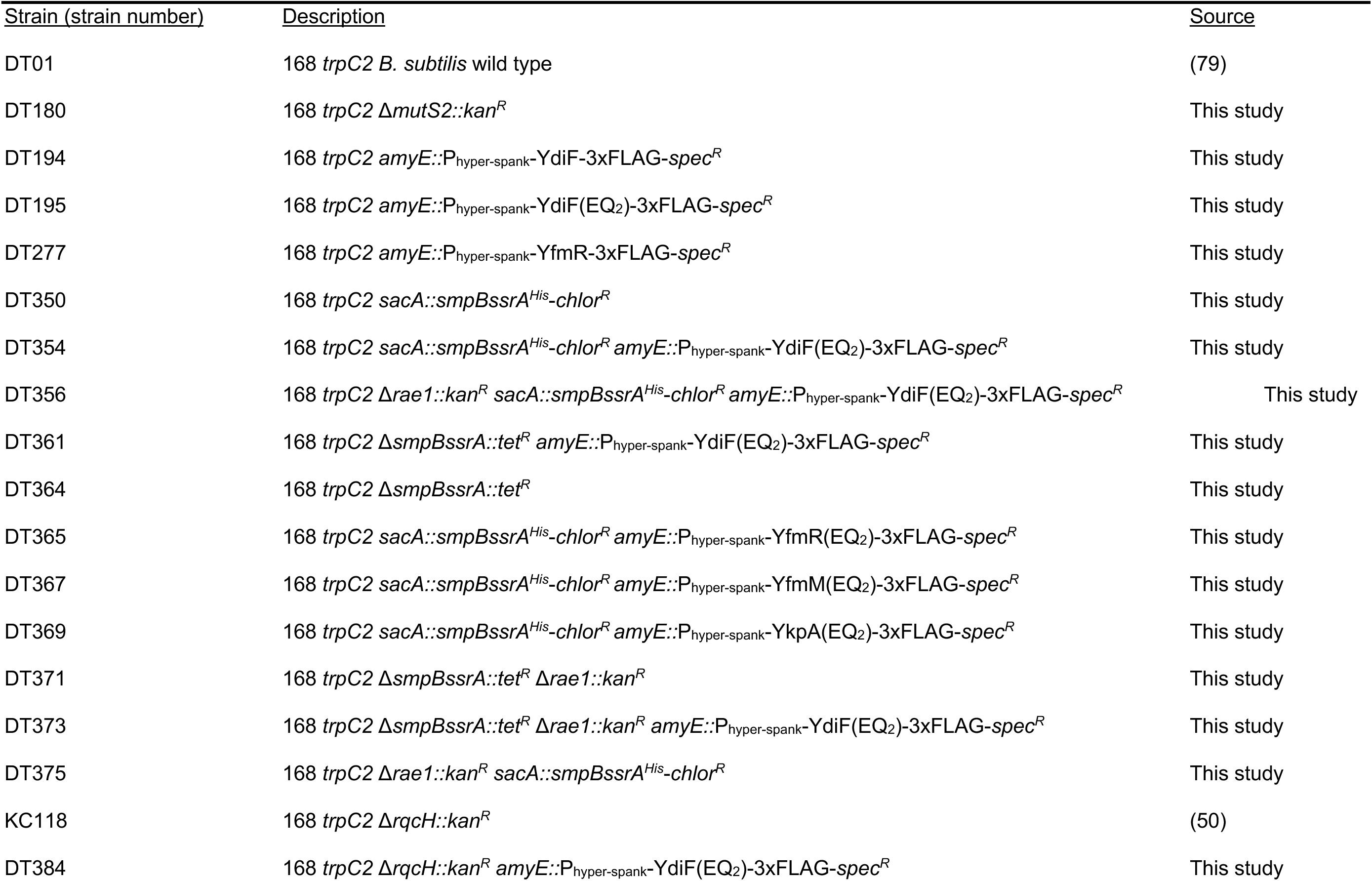
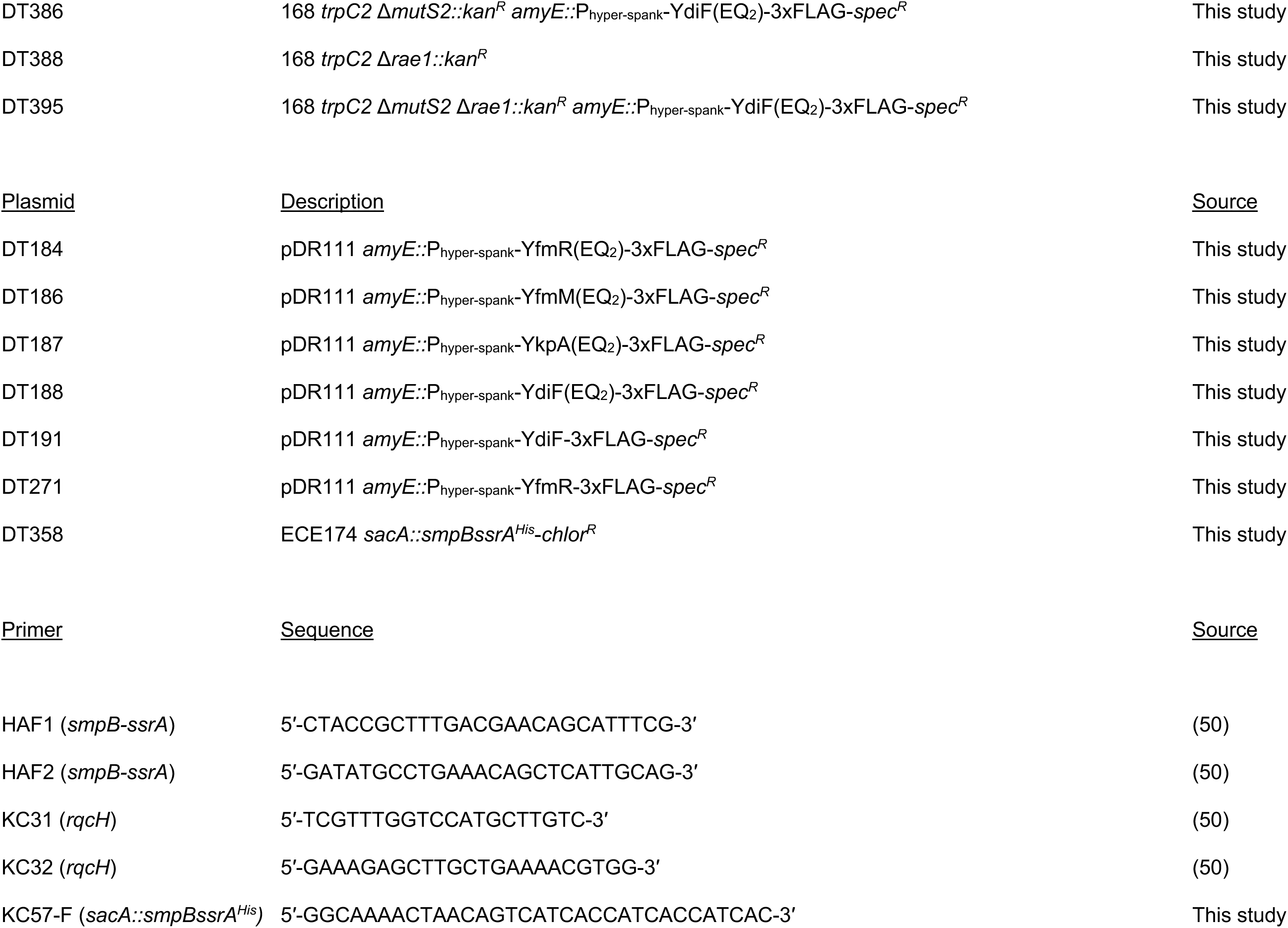
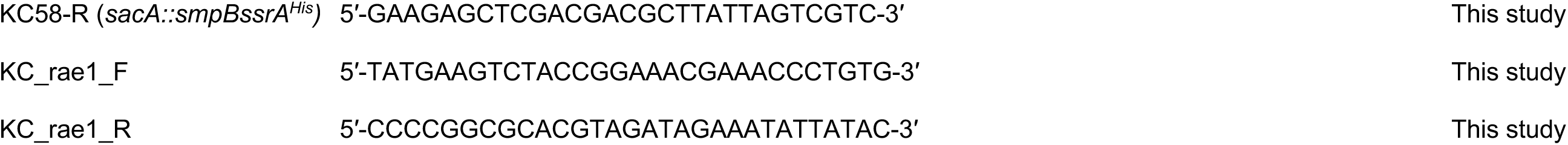

## Supplemental Figures

**Fig. S1.**
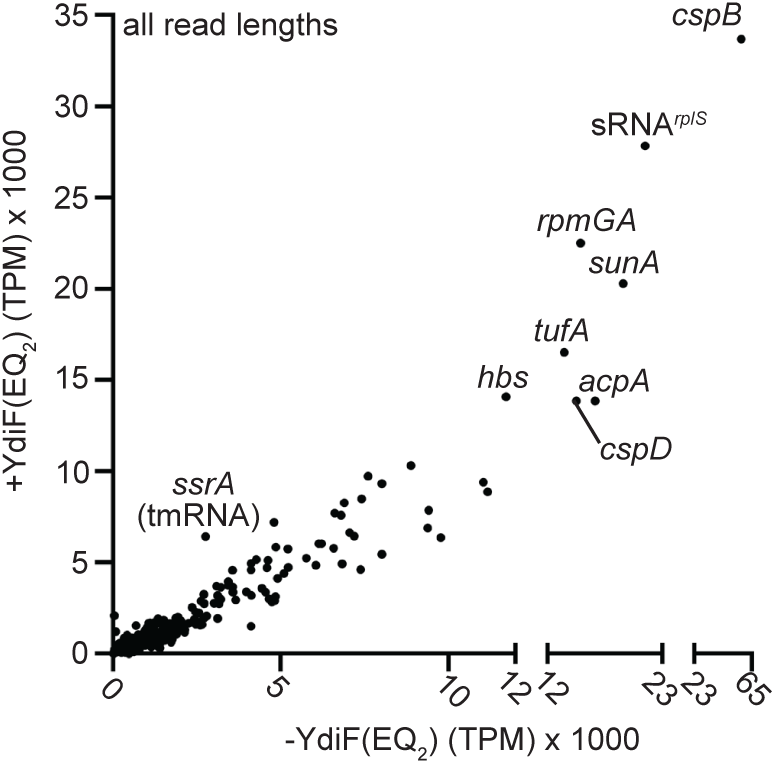
Ribosome collisions generated by YdiF(EQ_2_) are enriched for tmRNA. Scatterplot showing the average TPM of RPFs in wild-type and +YdiF(EQ_2_) cells yielded by mapping RPFs of all lengths.

**Fig. S2.**
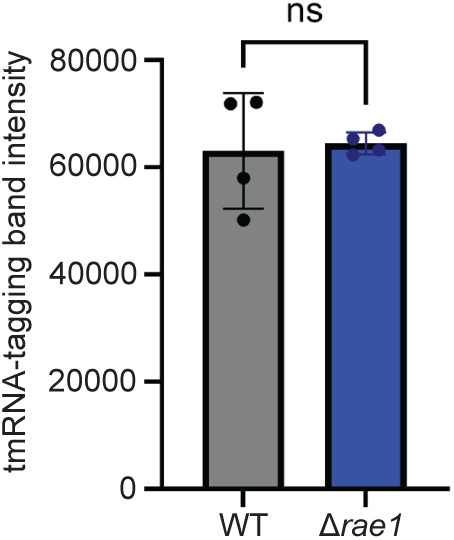
*rae1* deletion does not affect global *trans*-translation activity. Western band intensity quantification of *trans*-translation activity in tmRNA-His cells and in Δ*rae1* tmRNA-His cells. Cells were grown at 37°C in LB and harvested at OD600=1.0 before lysis for SDS-PAGE. Error bars represent the SD of biological quadricate tmRNA-His strains. The *p*-value is the result of Welch’s two-tailed T-test.

**Fig. S3.**
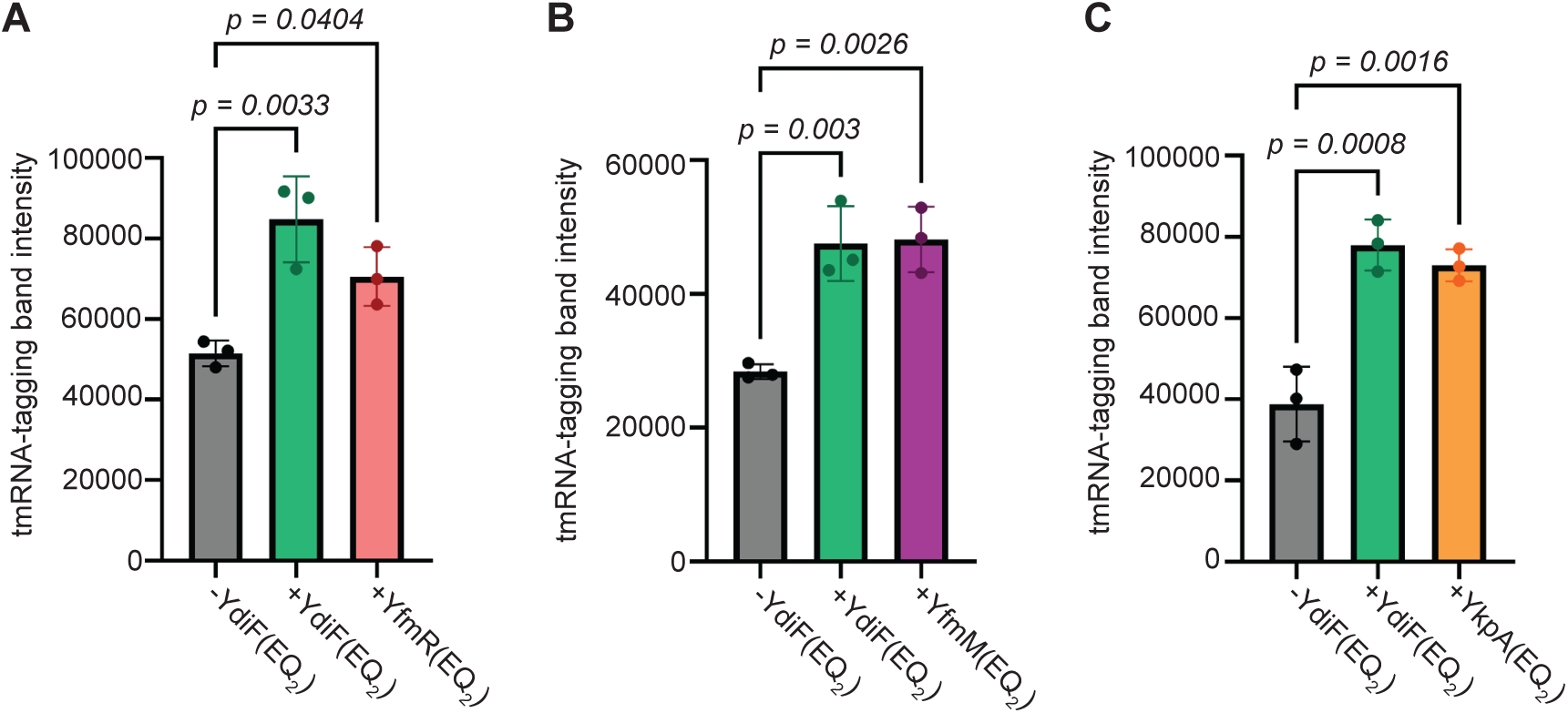
Expression of other ABCF ATPase(EQ_2_) mutants increases *trans*-translation activity. **A-C**, Western band intensity quantification of *trans*-translation activity in wild-type, +YdiF(EQ_2_), (**A**) +YfmR(EQ_2_), (**B**) +YfmM(EQ_2_), and (**C**) +YkpA(EQ_2_) cells. Cells were grown at 37°C in LB containing 2mM IPTG and harvested at OD600=1.0 before lysis for SDS-PAGE. Error bars represent the SD of biological triplicate tmRNA-His strains. *p*-values are the result of one-way ANOVA and Dunnett’s multiple comparisons test.

